# White matter microstructural integrity across the adult lifespan: Combined perspective of diffusion tensor and kurtosis imaging

**DOI:** 10.1101/2021.10.27.466088

**Authors:** Hiba Taha, Jordan A. Chad, J. Jean Chen

**Author notes:** Corresponding author: Jean Chen.

## Abstract

Studies of healthy brain aging have reported diffusivity patterns associated with white matter degeneration using diffusion tensor imaging (DTI), which assumes that diffusion measured at the typical b-value (approximately 1000 s/mm^2^) is Gaussian. Diffusion kurtosis imaging (DKI) is an extension of DTI that measures non-Gaussian diffusion (kurtosis) to better capture microenvironmental changes by incorporating additional data at a higher b-value. In this study, using UK Biobank data (b values of 1000 and 2000 s/mm^2^), we investigate (1) the extent of novel information gained from adding diffusional kurtosis to diffusivity observations in aging, and (2) how conventional DTI metrics in aging compare with diffusivity metrics derived from DKI, which are corrected for kurtosis. We find a general pattern of lower kurtosis alongside higher diffusivity among older adults. We also find differences between diffusivity metrics derived from DTI and DKI, emphasizing the importance of accounting for non-Gaussian diffusion. This work highlights the utility of measuring diffusional kurtosis as a simple addition to conventional diffusion imaging of aging.

## Introduction

An understanding of the diverse characteristics of brain degeneration in aging is needed for establishing benchmarks of healthy aging and detecting early signs of abnormal aging. Studies of healthy brain aging have previously reported patterns of white matter tissue and tract degeneration using diffusion tensor imaging (DTI). DTI parameters are derived from a 3-dimensional ellipsoid tensor model, which can be fit from diffusion MRI data acquired with a single diffusion gradient strength (typically b = 1000 s/mm^2^). The metrics derived from the DTI model include fractional anisotropy (FA) and mean diffusivity (MD), which is further broken down into axial diffusivity (AD) and radial diffusivity (RD) (Koerte and Muehlmann, 2014). DTI studies revealed age-associated decreases in fractional anisotropy (FA) and increases in mean diffusivity (MD), with RD increasing more than AD, suggesting an overall decrease in white matter integrity with advancing age (Bennett et al., 2009; Hsu et al., 2008; Hugenschmidt et al., 2008). However, conventional DTI assumes that diffusion in all direction is Gaussian, and thus fails to consider non-Gaussian diffusion in the white matter that may arise due to the presence of cellular membranes, multiple tissue compartments or metabolic debris, leading to limited specificity to microstructure. Without properly accounting for non-Gaussian diffusivity changes, how white matter tissue complexity changes with age is less understood.

Diffusion kurtosis imaging (DKI) is an extension of DTI that can better capture these microenvironmental changes (Jensen et al., 2005). In contrast to the Gaussian diffusion modelling of DTI, DKI measures the kurtosis of water diffusivity, which reflects the physical presence of biological structures and heterogeneity. In other words, kurtosis provides a measure of the restriction of Gaussian movement and can be used as a marker of tissue complexity. Higher kurtosis values can be interpreted to reflect the presence of biological barriers that conventional DTI fails to capture. DKI typically requires additional diffusion MRI data acquired with diffusion gradient strengths at higher b values, one at least 2000 s/mm^2^, since non-Gaussian effects are more prevalent at higher b-values whereas they are assumed to be negligible at the typical b-values used for DTI. DKI provides metrics such as axial kurtosis (AK), radial kurtosis (RK), and mean kurtosis (MK), and can also provide Gaussian diffusion metric estimates that are corrected for kurtosis effects (Steven et al., 2014).

Very few studies of brain aging have used DKI. In research published to date, reductions in kurtosis metrics MK, AK, and RK with age have been observed, with RK decreasing more than AK. These findings suggest overall reductions in diffusional heterogeneity, and therefore, decreases in white matter tissue complexity with age (Coutu et al., 2014; Das et al., 2017; Falangola et al., 2008; Lätt et al., 2013). Existing DKI literature thus establishes the existence of two types of microstructural variations with advancing age -- increasing diffusivity and decreasing diffusional kurtosis. Coutu et al. compared findings based on kurtosis and diffusivity (Coutu et al., 2014). It was shown that regions demonstrating similar diffusivity and kurtosis co-variations clustered together such that different clusters were associated with different spatial distributions of WM changes in aging. Subsequently, Billiet et al. demonstrated covariations between DKI and myelin-water fraction measurements in aging. RD and MK were found to covary in aging, but kurtosis age effects were minimal in the frontal lobe, where the strongest diffusivity-age associations are found (Billiet et al., 2015). The Coutu and Billiet studies differ in their report of region-specific associations between DKI and DTI associations with age, so a better understanding of the utility of combining DKI and DTI in studying aging is needed. Furthermore, in these studies DTI metrics were obtained from conventional DTI modeling rather than the DKI model, and it is currently unclear how significantly conventional DTI metrics are biased by non-Gaussian effects, since DTI metrics can only be corrected for kurtosis through DKI modeling.

In this study, we aim to address two main questions: (1) what information can be gained from adding DKI to DTI observations in aging? and (2) how do conventional and DKI-corrected DTI parameters differ in aging? To enhance the reliability of the results, we chose a sizable sample of 700 cross-sectional data sets (UKBiobank). We find interesting age associations in posterior-inferior brain regions that would not be fully interpretable without having access to both DTI and DKI results, and we find that DKI further allows for independent support of an interpretation of DTI age associations in crossing fiber regions.

## Materials and Methods

### Theory

The DTI and DKI models are given by the Einstein summations in Eq. (1) and (2), respectively,

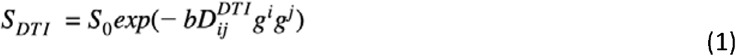

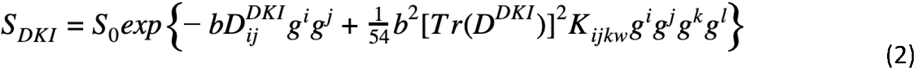

where S_0_ is the non-diffusion-weighted signal intensity, b and g are the diffusion gradient strength and direction, respectively, D is the diffusion tensor, K is the kurtosis tensor, and {i, j, k, l} are the summation indices. The diffusion tensors derived from the DTI and DKI models (D^DTI^ and D^DKI^ respectively) are not identical. By equating Eq. (1) and (2) along a given direction, we can see that age effects on the Gaussian diffusion coefficient *D*_DTI_ relate to age effects on the kurtosis-corrected diffusion coefficient *D*_DKI_ and kurtosis coefficient *K*, as well as the b-value *b*, by

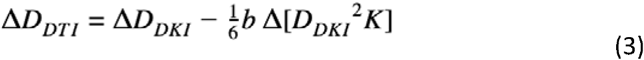

We see that while the discrepancy between *D*_DTI_ and *D*_DKI_ is larger at higher b-values, the two diffusion coefficients are never identical, even at low b-values, indicating that conventional DTI measurements acquired at the typical DTI b-value of 1000 s/mm^2^ are partly biased by kurtosis.

### Study Participants

All data sets used in this study were taken from participants of the UK Biobank Initiative. The UK Biobank *(ukbiobank.co.uk)* recruited over 500,000 participants aged 40-69 across the United Kingdom between 2006 and 2010, with the goal of imaging 100,000 participants. The UK Biobank received ethical approval under Research Ethics Committee reference number 16/NW/0274. All participants provided written consent.

The current study was conducted as part of the UK Biobank Application 40922. 22,427 participants aged 44 to 80 years were imaged at the time of the application. Participants reporting neurological, psychological or psychiatric orders, neurological injury or history of stroke were excluded from this study. The final group of participants consisted of a total of 700 healthy adults (50% male, 50% female) aged 46-80 years. This dataset has previously been reported on using the conventional DTI model (Chad et al., 2021).

### Diffusion MRI data

Diffusion MRI data were acquired using a Siemens Skyra 3T system with 5 b=0, 50 b=1000s/mm^2^, and 50 b=2000s/mm^2^ across 100 distinct directions at TR = 3.6s, TE = 92ms, matrix size = 104×104×72 with 8mm^3^ resolution, 6/8 partial Fourier, 3x multi-slice acceleration and no in-plane acceleration. Additional 3 b=0 volumes were attained with reversed phase encoding.

### Data Processing

Data was corrected for eddy currents and susceptibility-related distortions using EDDY and TOPUP, respectively. Additional details regarding the MRI acquisition and processing is provided by Alfaro-Almagro et al. (2018). Diffusion tensor and kurtosis models were fitted using the standard kurtosis-fitting approach implemented in the Diffusion Kurtosis Imaging Matlab Toolbox (https://cai2r.net/resources/software/diffusion-kurtosis-imaging-matlab-toolbox)(Veraart et al., 2013). Based on weighted linear least-squares estimation (Veraart et al., 2013), fractional anisotropy (FA), axial diffusivity (AD), mean diffusivity (MD), radial diffusivity (RD) and mean kurtosis (MK), axial kurtosis (AK), and radial kurtosis (RK) were obtained (Jensen and Helpern, 2010). AK maps were obtained by estimating the apparent diffusional kurtosis in the principal diffusion direction, while RK maps were obtained by averaging around all directions perpendicular to the principal diffusion direction. Note that the DTI parameters are obtained in two ways, using (1) the DKI model, which involves 2-shell data; (2) conventional DTI, which uses only the b = 1000 shell (pre-processed by the UKBiobank, as per Alfaro-Almargo et al., 2018). An additional constriction on kurtosis values (limiting values to be between 0 and 5) was required to increase the quality of derived maps and remove noisy voxels.

### Statistical Analysis

To assess whole-brain white matter age-related associations of these metrics, diffusion and kurtosis metric maps of each participant were registered to a mean FA skeleton in standard MNI152 space using FSL’s Tract-Based Spatial Statistics (TBSS) to obtain a WM skeleton. The mean FA skeleton was created with a threshold value of 0.2. Each map was then subjected to significance testing using FSL randomise with 500 permutations using threshold-free cluster enhancement (TFCE). A significant correlation of metric with age was defined by p < 0.05. Significant age-related metric associations per year were extracted and controlled for sex using a general linear model (mri_glmfit) as implemented in FreeSurfer. Neuroanatomical structures were identified using FSL’s integrated JHU White-Matter Tractography Atlas and the JHU ICBM-DTI-81 White Matter Labels.

To further study DTI and DKI linear age effects simultaneously, overlapping maps of different metric age effects including AK, RK, FA, AD and RD were derived using Matlab. Maps are displayed with the mean FA and mean FA skeleton derived from FSL’s TBSS statistical analysis procedure.

Additionally, a series of linear regression analyses were performed to assess DTI and DKI metric changes with age throughout the WM skeleton. Significant DTI and DKI metric values were derived using FSL’s *fslmeants* tool from the averaged DTI or DKI skeleton obtained from TBSS using a p-value of <0.05 and plotted with age using Matlab.

## Results

Significant global linear age effects on kurtosis are shown in **Fig. 1**, as represented by AK, MK and RK throughout the white matter skeleton. MK age effects were stronger in frontal and lateral regions including the forceps minor, anterior thalamic radiations, anterior corona radiata, and the cingulum compared to posterior regions (**Fig. 1A**). AK exhibited the weakest age effects compared to MK and RK (**Fig. 1B**). The strongest AK age effects were captured in the left and right superior, anterior and posterior corona radiata, and some portions of the left and right inferior fronto-occipital fasciculi. Posterior regions of the white matter skeleton within the occipital lobe exhibited little-to-no AK effects. Unilateral positive age associations of AK were also captured along the left body and genu of the corpus callosum and the left portions of the forceps minor, but not the right (see **Fig. 1B**). RK age effects were the strongest out of the investigated DKI metrics, spanning throughout the white matter skeleton with the exception of the posterior limb of the left and right internal capsule and portions of the forceps minor and forceps major (**Fig. 1C**). Some regions that did not exhibit significant AK effects reflected strong RK age effects, such as in the posterior regions of the white matter skeleton.

**Figure 1.**
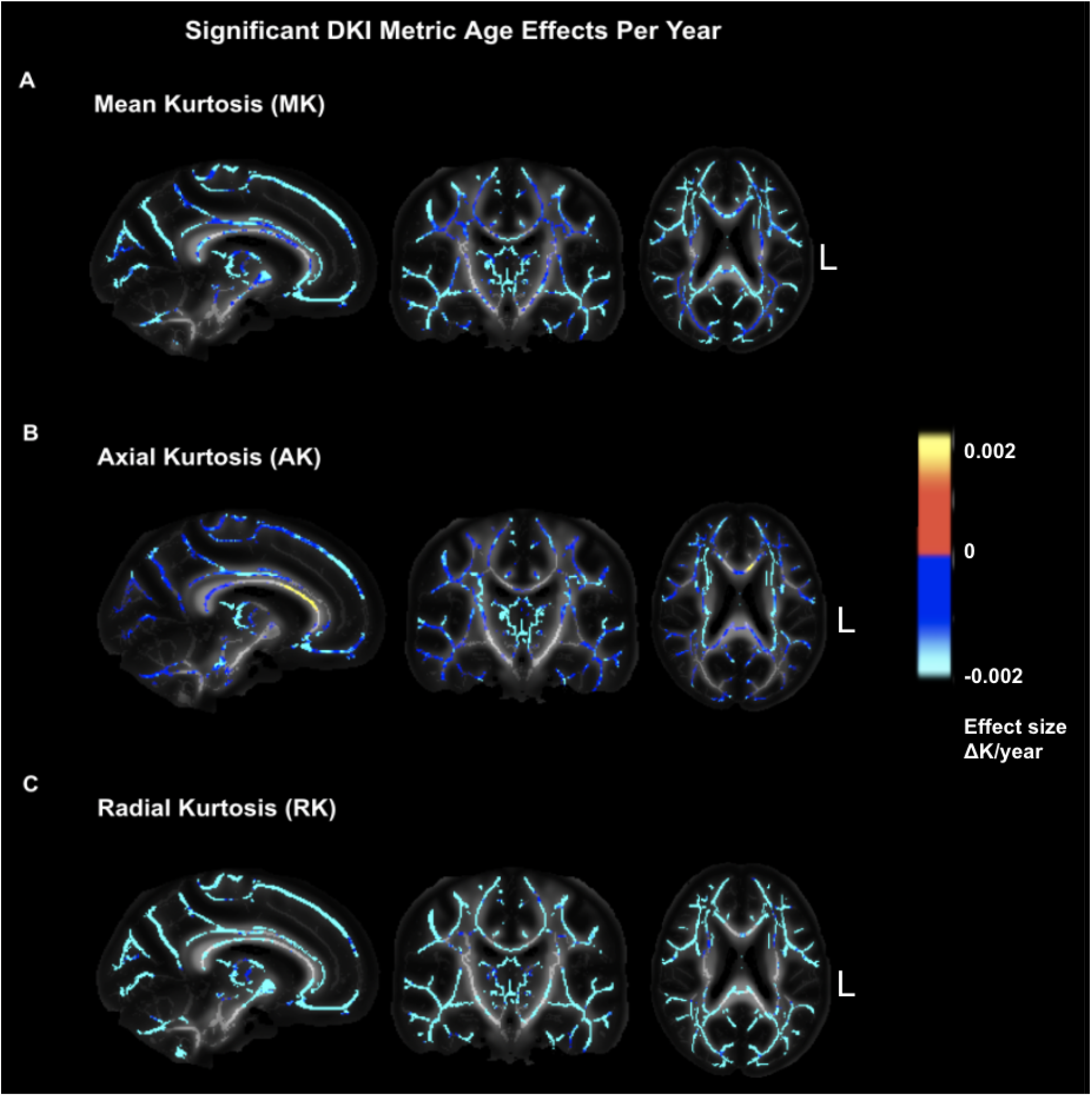
Age effects on DKI metrics across the WM skeleton, for (A) mean kurtosis (MK), (B) axial kurtosis (AK) and (C) radial kurtosis (RK). All effect sizes are thresholded at p< 0.05 (corrected for multiple comparisons). Warm colours (red/yellow) represent positive age associations and cool colours (blue) show negative age associations, and the colour bars indicate the effect sizes. Voxels of the WM skeleton without significant kurtosis age effects are represented in white, and ‘L’ indicates left. Significant negative age associations are observed throughout the WM skeleton in all kurtosis metrics.

Gaussian diffusion age effects as represented by FA, MD, AD and RD are shown in **Fig. 2**. These parameters are obtained from the DKI model. General trends of positive age associations of diffusivity and negative age associations of FA were captured throughout the white matter skeleton. Additionally, significant positive age associations of FA were captured in the right and left corticospinal regions, including the superior and posterior corona radiata, superior portions of the posterior limb of the internal capsule and the retrolenticular parts of the internal capsule, with larger age effects observed on the right regional structures (**Fig. 2A**). Positive age associations of MD were widespread throughout the white matter skeleton, with the exception of some posterior regions and inferior regions in which no significant MD effects were observed, including the posterior limb of the internal capsule and the cerebral peduncles (**Fig. 2B**). AD exhibited a general trend of positive age effects in frontal and superior regions, including the anterior limb of the internal capsule, and negative age effects in posterior and inferior regions of the white matter skeleton, including the posterior limb of the internal capsule (**Fig. 2C**). Similar to MD, RD positive age effects were observed largely across the white matter skeleton (**Fig. 2D**). Posterior regions with negative AD age effects including the posterior limb of the internal capsule notably exhibited overlapping RD positive age effects and FA negative age effects.

**Figure 2.**
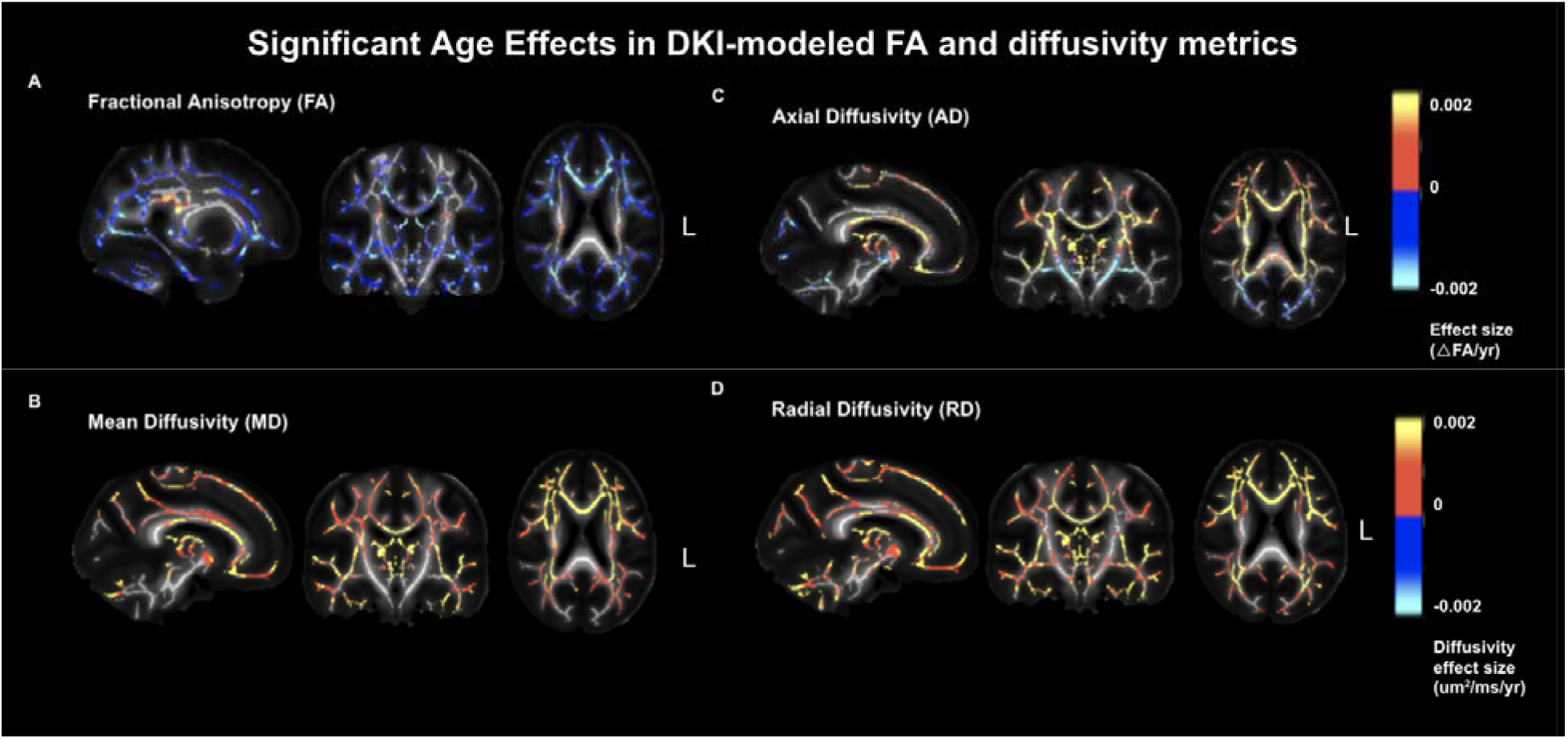
Age effects on DKI-modeled DTI metrics across the WM skeleton, for (A) fractional anisotropy (FA) [ΔFA/year], (B) mean diffusivity (MD)[μm/ms^2^/year], (C) axial diffusivity [μm/ms^2^/year] and (D) radial diffusivity [μm/ms^2^/year]. All effect sizes are thresholded at p< 0.05 (corrected for multiple comparisons). Warm colours (red/yellow) represent positive age associations and cool colours (blue) show negative age associations. Brighter colours represent greater effect sizes. Voxels of the WM skeleton without significant kurtosis age effects are represented in white, and ‘L’ indicates left. Diffusivity metrics mostly exhibited significant positive age associations (p < 0.05), whereas FA showed negative age associations throughout the WM skeleton. Positive age associations of FA were captured in the left and right corticospinal regions. Additionally, positive age associations of AD were observed in superior and frontal regions while specific negative age associations were observed in posterior and inferior regions.

To provide a more comprehensive report on white matter microstructural changes with age, DTI and DKI age effects were explored simultaneously by investigating a series of overlapping effects of different combinations of DTI and DKI metrics shown in **Fig. 3**. A summary of age associations in various metrics within various major WM pathways is provided in **Table S1** in Supplementary Materials. First, it can be observed (from **Fig. 3A**) that MK and MD age effects overlap (in opposite directions) throughout nearly the entire white matter skeleton, with the exception of posterior-inferior white matter, where significant MK reduction but no significant MD increase with age is observed (indicated in blue). Moreover, negative AK age effects overlap with positive AD age effects in the majority of anterior-superior white matter, including the internal-capsule region, but AD does not display positive age effects in posterior-inferior regions (**Fig. 3B**). It is shown in **Fig. 3C** that AD exhibits negative instead of positive age associations in these regions. The negative AD-age associations overlap with positive RD-age associations. AK is negatively associated with age through most of this region, although in some areas negative RK age effects are observed without concurrent AK negative age effects (**Fig. 3D**). As well, in the corona radiata, only negative AK-age associations are seen, without concurrent RK effects. Positive RD age associations generally overlap with negative RK age effects across the white matter, except for in parts of the anterior corpus callosum and posterior white matter (**Fig. 3E**). Positive FA age associations overlap with negative AK associations in parts of the corona radiata alone (**Fig. 3F**), but in the remainder of the white matter, negative age associations in FA are observed, and they do not always overlap with AK effects (**Fig. 3G**). Lastly, the spatial distributions of negative age effects in both AK and AD is illustrated, with the overlap in the posterior-inferior tracts only (**Fig. 3H**).

**Figure 3.**
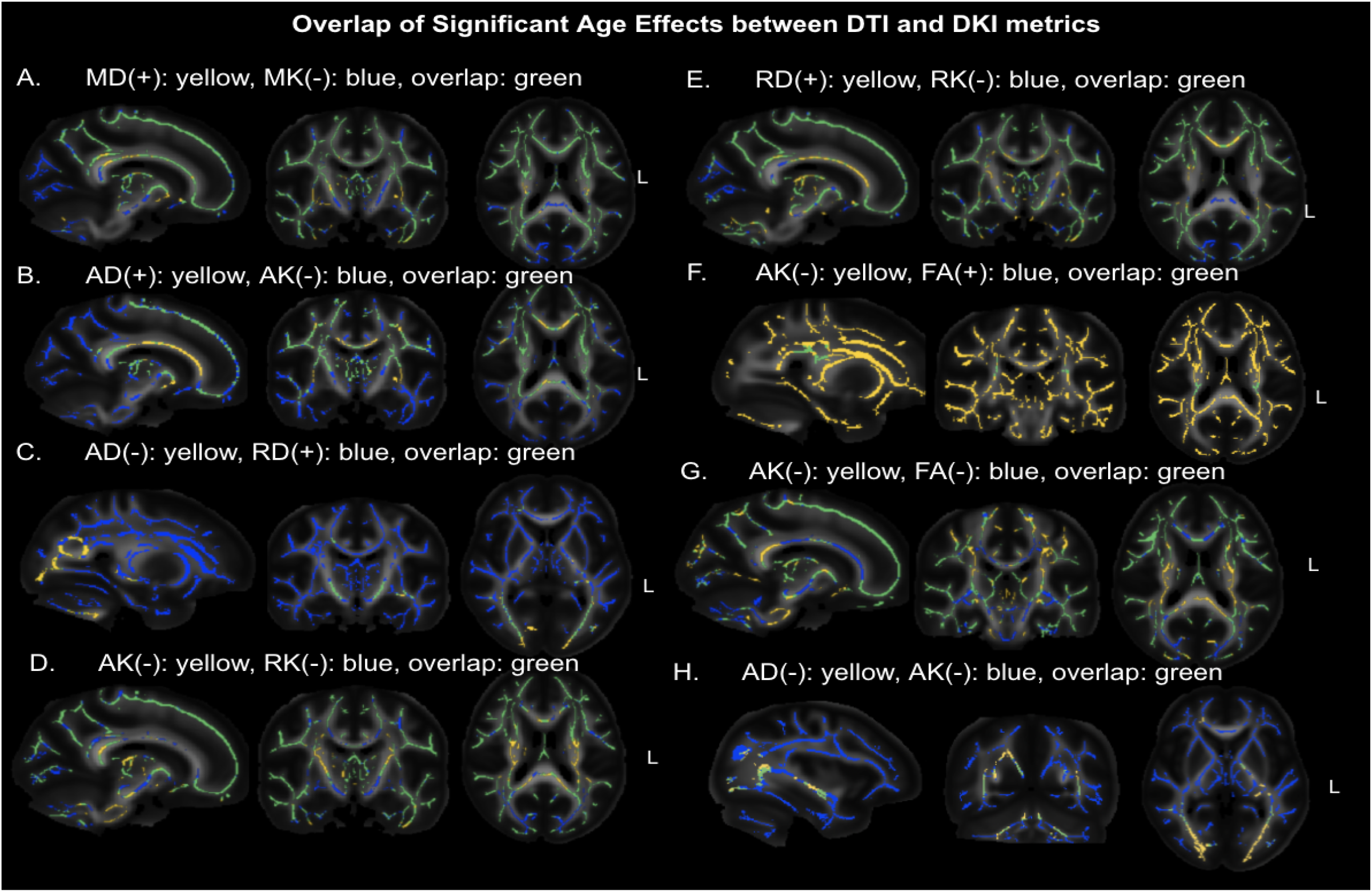
Overlap between significant DKI-modeled DTI and DKI age effects (p < 0.05) across the WM skeleton. All images are overlaid on the mean FA map, and ‘L’ indicates left, (−) indicates significant negative association with age, while (+) indicates significant positive association with age.

Shown in **Fig. 4** are scatter plots for regions of overlapping influence. The region associated with **Fig. 4A** can be gleaned from **Fig. 3B** and **3E**, in which AD and RD display overlapping positive associations with age across the entire white matter, with the exception of the posterior-inferior region and the areas where FA is positively associated with age. **Fig. 4B** is plotted from the region of negative AD and positive RD age associations in **Fig. 3C**. Finally, AK and RK are both negatively associated with age in much of the white matter and plotted in **Fig. 4C**.

**Figure 4.**
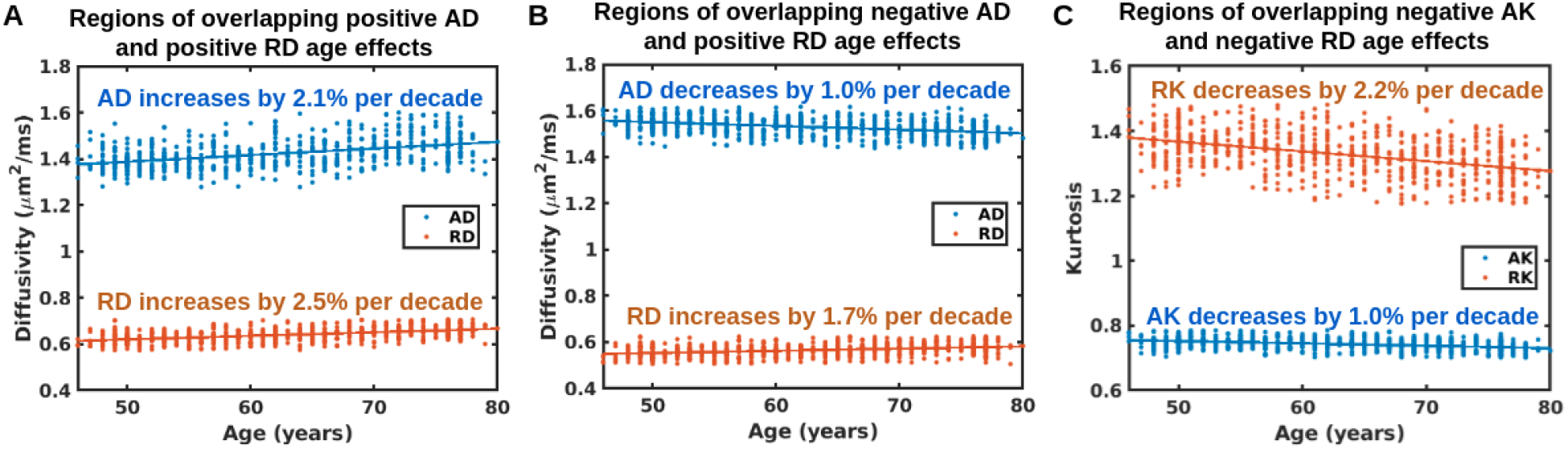
Comparisons of DKI-modeled axial/radial diffusivity and kurtosis associations with age, in (A) regions of overlapping positive AD and RD age effects, (B) regions of overlapping but opposite AD (negative) and RD (positive) age effects and (C) regions of overlapping negative AK and RK age effects. The red lines represent the best linear fits.

Summarized in **Fig. 5** is a direct comparison of DTI parameters FA and MD as derived from the DKI and DTI models. FA derived from the DKI model exhibits greater negative age effects and smaller positive age effects than FA derived from the DTI model throughout most of the white matter (**Fig. 5A**), whereas MD derived from the DKI model exhibits greater positive age effects than MD derived from the DTI mode throughout most of the white matter (**Fig. 5B**). DKI provides higher FA estimates than DTI (compare **Fig. 5C** and **Fig. 5D**), and DKI provides higher MD estimated than DTI (**Fig. 5E**).

**Figure 5.**
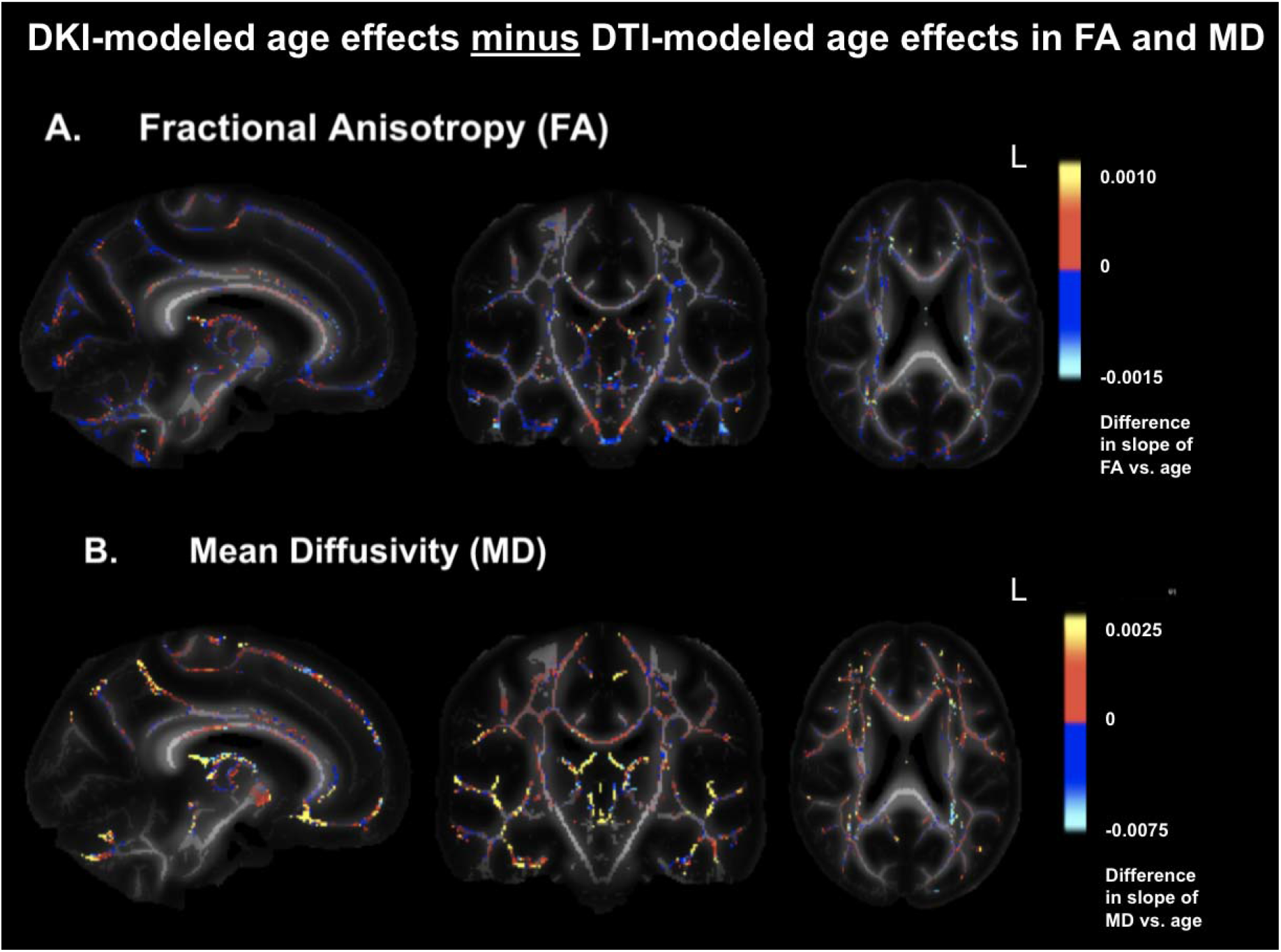
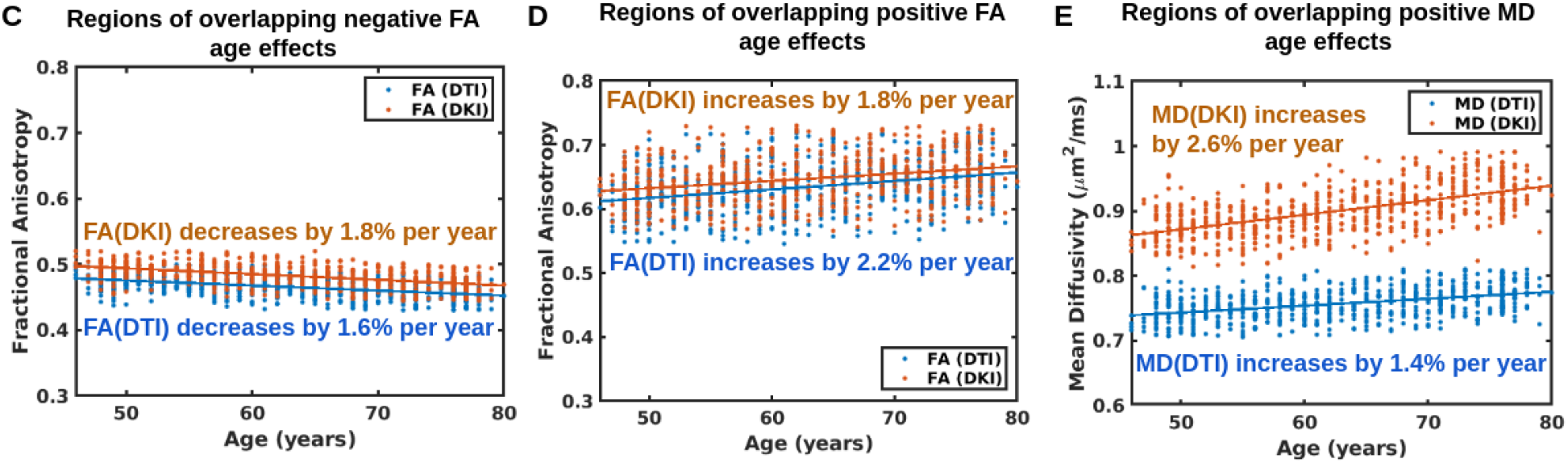
MD and FA estimated with DTI and DKI models. All differences are shown only for age effects thresholded at p< 0.05 (corrected for multiple comparisons). (A) DKI-corrected FA displays weaker age associations; (B) DKI-corrected MD displays stronger age associations. Scatter plots corresponding to these comparisons are also shown for (C) negative FA associations with age, (D) positive FA associations with age and (E) MD associations with age. The red lines represent the best linear fits, and ‘L’ indicates neurological left.

Illustrated in **Fig. 6** is a dissection of diffusivity age effects estimated using the DKI and DTI models. Kurtosis correction reveals negative age associations in the posterior WM tracts that cannot be observed with conventional AD **(Fig. 6a & 6B)**. Conversely, DKI and DTI estimated RD exhibit very similar age effects **(Fig. 6C & 6D)**.

**Figure 6.**
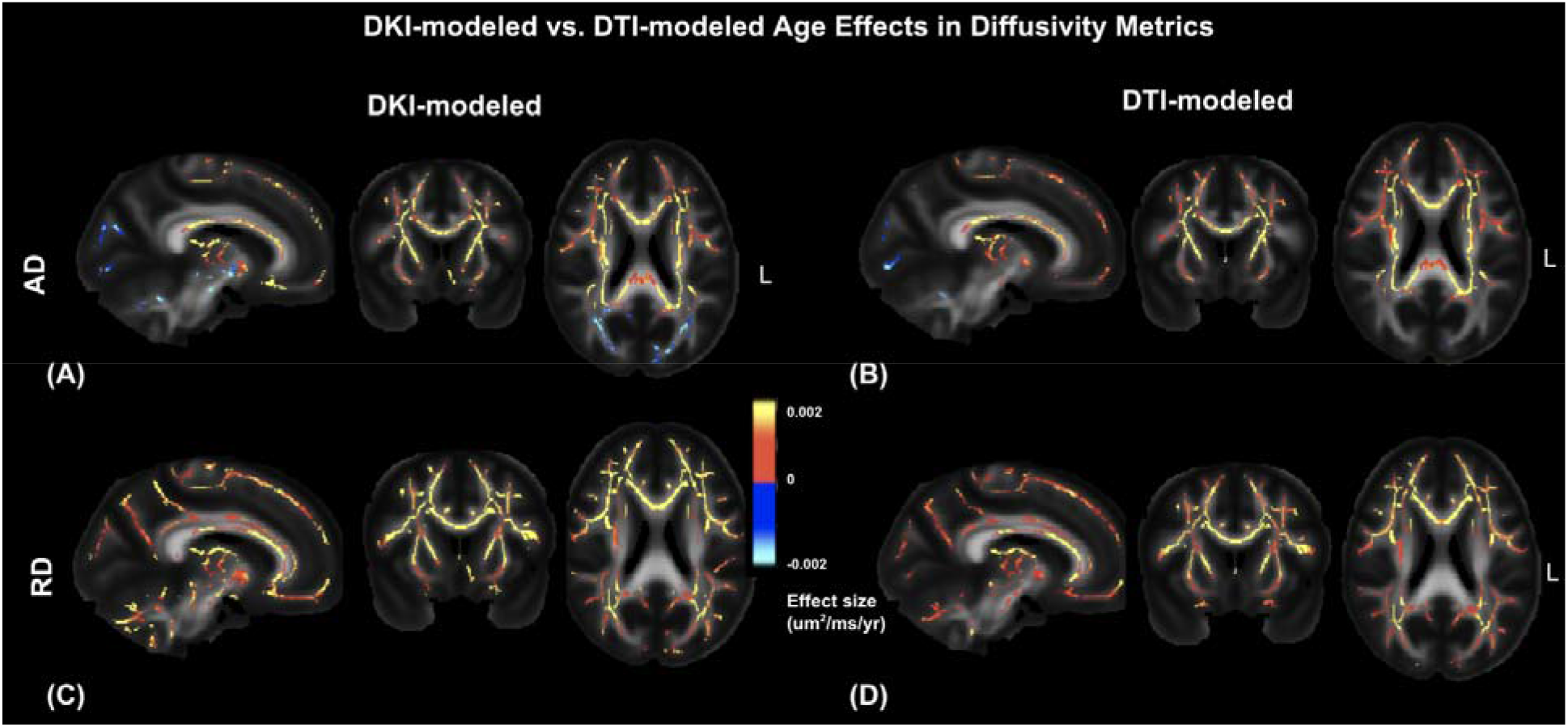
Diffusivity age effects estimated with DKI and DTI models. All effect sizes are thresholded at p< 0.05 (corrected for multiple comparisons). DKI-corrected AD (A) displays more pronounced negative age associations than DTI-derived AD (B). For RD, the difference between DKI-modeled (C) and DTI-modeled (D) age effects is less pronounced. ‘L’ indicates neurological left, and the colour bar indicates effect size.

## Discussion

DKI offers the value of quantifying white matter tissue complexity and compartmentalization beyond the capability of conventional DTI. Reduced kurtosis directly implies reduced non-Gaussianity in water movement and has been interpreted as signifying a reduction in microstructural heterogeneity. This interpretation has been histologically supported (Maiter et al., 2021) and also supported by multi-modal MRI studies of aging, where DKI effects have been found to be distinct from diffusivity effects: In a study of 59 participants, DKI metrics were shown to be distinctly correlated with myelin-water fraction (derived from multi-exponential T2 relaxation), and intracellular and extracellular water fractions (derived from NODDI) (Billiet et al., 2015). In the current study, we specifically report on the concomitance of kurtosis increases and diffusivity decreases with advancing age. In the context of combining the perspectives of DTI and DKI, we demonstrate the unique utility of adding DKI to DTI measurements that warrants the acquisition of multi-shell diffusion MRI data. Furthermore, as the DKI model also produces diffusivity metrics, it provides the opportunity to observe differences between conventional and kurtosis-corrected DTI metrics.

### Kurtosis associations with age

As detailed earlier, in contrast with the abundance of conventional DTI studies, there have been only a few previous reports describing WM diffusional kurtosis reductions in normal aging. Findings by Falangola et al. (Falangola et al., 2008) and Das et al. (Das et al., 2017) demonstrate that kurtosis is negatively associated with age. Also, reduced MK has found to be associated with reduced cerebral blood flow and cognitive function (Tu et al., 2021; Yang et al., 2021), and has been demonstrated as a potential marker for early detection of amyloid pathology (Praet et al., 2018). However, published studies generally have small sample sizes and do not quantitatively examine DKI and DTI in conjunction. One exception is a study by Benitez et al. (Benitez et al., 2018), in which MK was found to inversely associate with MD.

In our previous study on 111 healthy adults aged 33-91 years (Coutu et al., 2014), DKI parameters were derived using 3-shell data (b = 700, 1400 and 2100 s/mm^2^, 24 directions per b value), while DTI parameters were derived from the b=700 shell (based on the DTI model). RK and MK were both more strongly associated with age in superior and anterior than parietal and temporal regions, with the inferior posterior WM regions showing the least degree of association. These previous findings are largely supported by the results of the current study, performed on a much larger data set, although there are 2 major acquisition and analysis differences between the current approach and that which was employed in the Coutu paper: (1) our DKI metrics are derived using 2 shells (b = 1000 and 2000 s/mm2); (2) our DTI metrics are derived using both shells (through the DKI model).

We found a greater extent of WM showing kurtosis than diffusivity age effects **(compare Fig. 1 & 2)**. Notably, we observed an increase in AK in the body of the corpus callosum with age **(Fig. 1B and Fig. S1 (Supplementary Materials))**. This observation has previously been reported by Billiet et al. in a smaller cohort (Billiet et al., 2015), indicating increased tissue heterogeneity in the axial direction, and may derive from atrophy leading to increased partial voluming between tissue types in aging (as age effects on AK are typically lower than RK, an increase in kurtosis derived from partial voluming is expected to influence AK age effects more than RK). Moreover, the coexistence of increased AD with the increase in AK in this region **(Supplementary Figure S1)** supports the theory of altered partial-volume effects in this region, since increased AK is generally associated with decreased AD in single compartments. This was not observed in previous smaller studies (Falangola et al., 2008; Das et al., 2017), highlighting the power of a larger sample.

### Conventional DTI vs. kurtosis-corrected DTI

We note that the age effects on RK and kurtosis-corrected RD display highly similar spatial patterns **(Fig. 3C)**; since age effects are more pronounced radially than axially, the correspondence between RD and RK effects likely drives the similarities between MD and MK (compare **Fig. 1A and 2B**). We further categorized the DTI and DKI parameters by their collective trends **(Fig. 4)**.

An important novel finding is that in **Fig. 4B**, we show the scatter plots in regions of AD decrease overlapped with RD increase in aging, which is, as stated earlier, mainly found in the posterior-inferior WM **(Fig. 3C)**; here AD decreases at a rate of 1.0% per decade whereas RD increases at a rate of 1.7% per decade. Note that this is to our knowledge the first instance of age-associated decreases in DKI-corrected AD being reported, which is significant because this finding is in contrast with conventional DTI, whereby negligible negative age associations are observed in AD **(Fig. 6B)**. The finding of clear negative AD age associations after correcting for kurtosis is also in contrast to findings of Coutu et al., who reported only very weak positive AD-age associations in the posterior white matter, as well as to findings of Billet et al., who reported no significant AD-age associations in posterior tracts. Our findings show that by adapting a kurtosis-corrected diffusion model, the sensitivity to negative AD-age associations is improved.

A key message of this study is that there are two major (directionally-independent) processes happening in aging: changing diffusivities and changing kurtoses. **Figure 6** implies that age effects in kurtosis slightly influence diffusivity measurements at a typical b-value of 1000 s/mm^2^. The effect of kurtosis on diffusion parameters is greater when parameters are estimated using data acquired with high b-values (Eq. 3). Importantly, if both diffusivity and kurtosis were decreasing, these processes can effectively cancel each other in conventional DTI (less Gaussian diffusivity + less non-Gaussian restrictions manifest as no significant change in conventional DTI-based diffusivity). Therefore, with the DKI model, by accounting for kurtosis separately, we can more easily uncover decreased axial diffusivity with advancing age. For the same reason, MD exhibits greater increases and FA exhibits greater decreases in aging after correcting for kurtosis (see **Fig. 5**), because in conventional DTI these effects are masked by increasing kurtosis, especially along the radial direction.

### Combining DKI and DTI

When examining DKI and DTI in conjunction, the Coutu study found a great deal of shared variance between MK and MD, as well as between RK and RD. Moreover, in both pairs, the kurtosis metric accounted for more age-related variations than the diffusivity metric, contrary to findings by Falangola et al. based on 24 subjects (Falangola et al., 2008). Additionally, the Coutu study suggested that multiparametric combinations of DKI and DTI metrics, referred to as “diffusion footprints”, could be more informative of specific patterns of microstructural aging. Specifically, k-means clustering was used to group regions demonstrating similar FA, diffusivity and kurtosis co-variations. The superior frontal tracts are characterized by strong age effects across all parameters, while the inferior-posterior tracts are characterized by strong age effects in all but AD, consistent with the differences between early- and late-myelinating tracts reported previously (Benitez et al., 2018).

In this work, we expand on the idea of observing DKI and DTI metric variations in combination, and we uncover regions in which the previously observed trends do not apply. Specifically, we have identified two interesting combinations of parametric associations with age: (1) the posterior-inferior regions, which were previously found to display low AD sensitivity to age (Billiet et al., 2015; Coutu et al., 2014); (2) parts of the corona radiata, which is characterized by known fibre crossings (Chad et al., 2018; Douaud et al., 2011).

While positive age associations in diffusivity and negative age associations in kurtosis overlap in most brain regions, the posterior-inferior region is an exception. We observed decreasing AD in this region despite decreasing AK with advancing age. In other words, tissue heterogeneity is in decline axially, but the Gaussian aspect of axial diffusion is also reduced. One scenario that can explain this finding is that axial diffusion is being impeded despite the presence of microstructural degeneration that hinders Gaussian diffusion. Sources of such impediments, as previously discussed in our work, could be intra-axonal debris accumulation that takes place in early stages of neurodegeneration (Wyss-Coray, 2016). Once the myelin sheaths and axonal membrane are breached by greater degrees of degeneration, presumably the intra-axonal debris will disperse and no longer impede diffusion. Thus, we propose that the lack of positive AD-age association combined with a negative AK-age association could be used as an *early* marker of neurodegeneration, in contrast to the AD increase that takes place in the superior-frontal regions that likely reflects later stages of degeneration. This proposal is consistent with the “anterior-posterior” and “superior-inferior” gradients of aging (Madden et al., 2012) which in turn is related to the “last-in-first-out” theory (Bender et al., 2016): the posterior and inferior regions, being closest to the brainstem, are the earliest to myelinate in development and latest to degenerate in adult aging. To further verify the extent of demyelination in the posterior-inferior region in normal aging, it would be helpful to integrate myelin-water mapping with a diffusion imaging protocol in future work.

Thus far, most of our interpretations of both DTI and DKI findings have been assuming a single-fibre scenario, but many voxels contain multiple fibre crossings. A higher FA is typically indicative of stronger microstructural integrity. In parts of the corona radiata, however, we have shown previously using both free-water imaging (Chad et al., 2018), tractography (Chad et al., 2021) and orthogonal-tensor decomposition (Chad et al., 2021) that conventional FA can become misleadingly elevated in older age due to selective degeneration of the secondary fibre population relative to the primary fibre. In the eyes of DKI, such selective degeneration would translate into a reduction in AK due to the removal of tissue heterogeneity in the axial direction. Is this indeed what we observe in these regions? The areas of strongest negative AK-age association **(Fig. 1B)** indeed overlap with the areas of positive FA-age association **(Fig. 2A)**.

The schematic diagram in **Fig. 7** is inspired by the observations from this study. A healthy single-fibre model (representative of young controls) is shown in **Fig. 7A**. The zoomed version illustrates intact myelin sheaths and axonal membrane, which encapsulate intra-axonal debris (e.g. microfibrils, neurofilaments, etc). **Fig. 7B** and **7F** are associated with more commonly observed patterns, whereby advanced degeneration leads to observable increased diffusivity and decreased kurtosis. **Fig. 7C** and **7G** are less commonly observed, but nonetheless are interesting cases that add valuable understanding to the degenerative process, as these DTI and DKI trends are observed in parts of the corona radiata in our current study. This schematic demonstrates that when studying white-matter degeneration, the combined use of DKI and DTI, i.e. combining metrics of diffusion anisotropy, diffusivity and kurtosis to form “diffusion footprints”, can aid in biological interpretation. For instance, in the scenario of **Fig. 7C**, in the earliest stage of neurodegeneration, diffusion is enhanced radially due to demyelination, but also impeded (most pronounced axially) due to the accumulation of intra-axonal debris escaping into extracellular space. Here, the utility of kurtosis metrics stem from their ability to distinguish between variability in extracellular hindrance of diffusion (such as due to myelin layers), which can be probed using Gaussian DTI metrics, and intracellular restriction of diffusion (such as due to metabolic damage and accumulated debris), for which kurtosis metrics are needed. Unlike complex tissue structures, debris is expected to be randomly distributed, resulting in a reduction in AK and RK. This interpretation is consistent with our observations in the posterior-inferior WM **(Fig. 3H)**, and applies to the equivalent scenario in cross-fibre regions (**Fig. 7G**, corresponding to **Fig. 3F**), with the exception of early selective degeneration leading to an increase (rather than a reduction) in AD and FA. This scenario, if continued, would result in the complete degeneration of the secondary fibre. We propose this set of schematics as a guide for the use of multi-parametric diffusion imaging to discover more subtle signs of early degeneration, particularly in regions of complex fibre architectures; however, we also emphasize that diffusion processes are complex and the simplistic schematics should always be used with caution.

**Figure 7.**
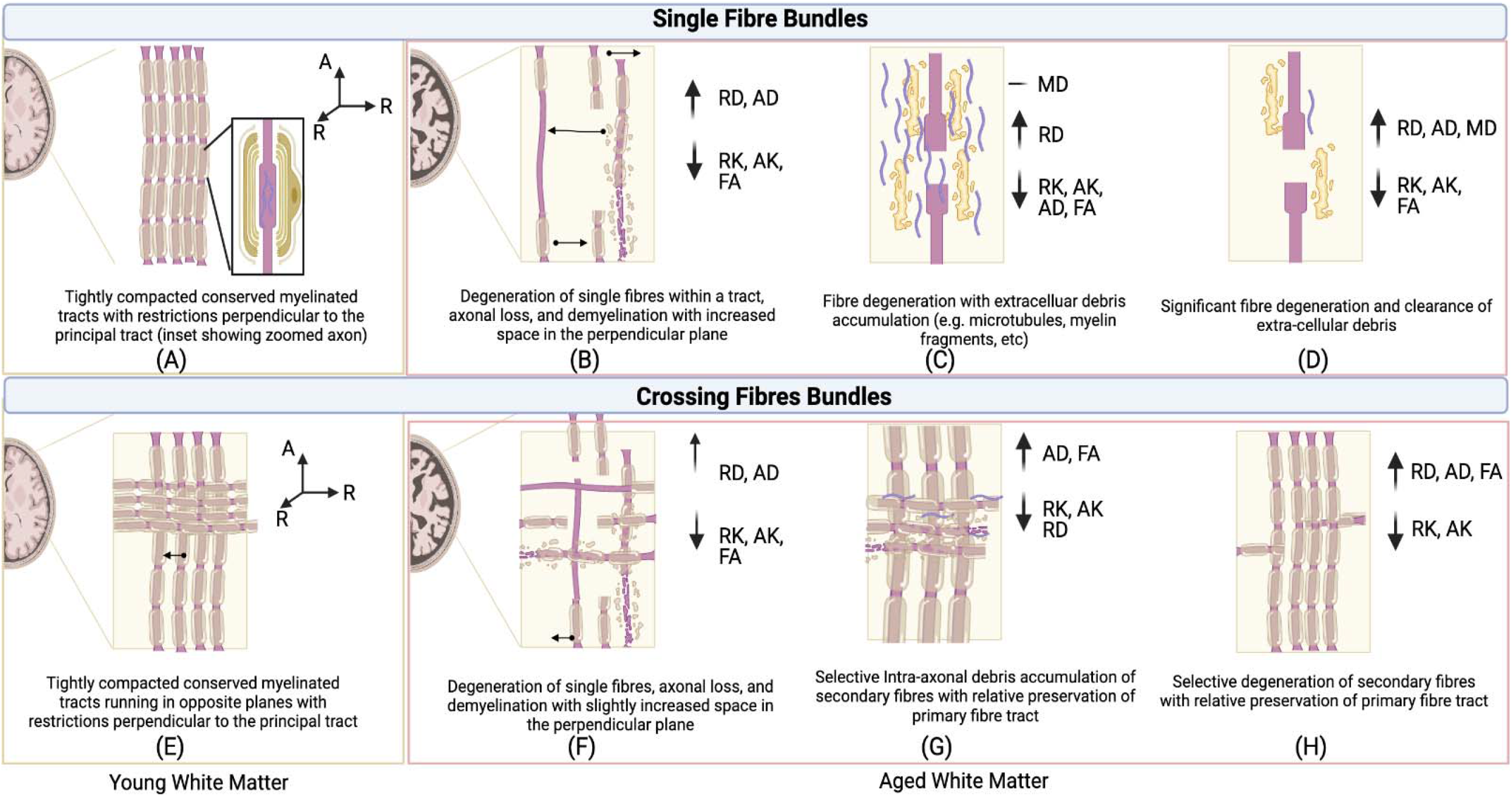
A schematic diagram summarizing the likely interpretations of various combinations of DTI and DKI parameters’ associations with age. (A) and (E) represent younger white matter in the single-fibre and crossing-fibre scenarios. (B-D) represent three different scenarios of single-fibre degeneration, while (F-H) represent three cases of crossing-fibre degeneration.

### Limitations and future work

There is currently no standard tool for quantifying kurtosis. In addition to the Matlab toolbox used in this work, we are also aware of others such as the DiPy toolbox (www.dipy.org) and the diffusion kurtosis estimator (DKE) (https://www.nitrc.org/projects/dke/). In the course of the analysis, we have discovered that the observed age effects on the DKI and simultaneously generated DTI metrics can differ between tools (Taha et al., 2021). In our future work, we plan to expand the comparison of toolboxes and involve the developers in the process, as the toolboxes are known to evolve rapidly.

In previous work, a quadratic AK-age association was investigated (Coutu et al., 2014). This was not pursued in our approach, as the data did not yield any noticeable non-linear trends. Moreover, as alluded to before, as myelin-water mapping is not available through the UKBiobank data set, we are unable to expand our validation of the hypothesis that “intra-axonal debris accumulation can lead to the observed reductions in AD and RK in aging”. In future work, the data can be fit to models like NODDI (Zhang et al., 2012), which provides metrics for cross-validation, such as fibre volume and dispersion. The question of interpretation could be investigated by a dedicated study as part of our future work.

## Acknowledgments

This work was funded by the Canadian Institute of Health Research and the Canada Research Chairs Program. We are also grateful for the research infrastructure provided by Compute Canada. We also thank Dr. David Salat for valuable feedback.

## Notes

### Competing Interest Statement

The authors have declared no competing interest.

