## Supplementary Materials for "White matter microstructural integrity across the adult lifespan: Combined perspective of diffusion tensor and kurtosis imaging"

**Table S1. Summary of age associations in various WM tracts**, as shown by DKI and DTI parameters.

| **Crossing fibre regions ↑↓** | **Single Fibre regions ↑↓** |
| --- | --- |
| **Corpus Callosum**   - **Splenium** [↑ AD, **↓** MK] - **Body** [**↓** FA, ↑ AD, ↑ RD, ↑ MD, **↓** AK, **↓** MK] - **↓** integrity, ↑ diffusion, **↓** kurtosis - **Genu** [↑ RD, **↓** AK, **↓** RK, **↓** MK] - ↑ radial diffusion, **↓** kurtosis   **Uncinate Fasciculus** [116, 138, 62]  [↑ AD, ↑ RD, ↑ MD, **↓** AK, **↓** RK, **↓** MK]   - ↑ diffusion, **↓** kurtosis   **pCorona Radiata** [64, 106, 93] (*superior* *portions of the CST*)  [**↑** FA, ↑ AD, ↑ MD, **↓** AK, **↓** RK, **↓** MK]   - ↑ anisotropy, ↑ diffusion, **↓** kurtosis   **Anterior limb of the internal capsule** [70, 106, 69]  [**↓** FA, ↑ AD, ↑ RD, ↑ MD, **↓** AK, ↓ RK]   - **↓** integrity, ↑ diffusion, **↓** axial and radial kurtosis   **Posterior limb of the internal capsule** [70, 106, 69] (*superior portions*)  [**↓** FA, **↓** AD, ↑ RD, ↑ MD]   - **↓** integrity, ↑ diffusion   **Posterior limb of the internal capsule** [70, 106, 69] (*inferior portions*)  [**↓** FA, **↓** AD, ↑ RD, ↑ MD, **↓** AK, ↓ RK, **↓** MK]   - **↓** integrity, ↑ radial diffusion, **↓** kurtosis   **Cingulum** [98, 152, 93]  [**↓** FA, ↑ RD, ↑ MD, **↓** AK, ↓ RK, **↓** MK]   - **↓** anisotropy, ↑ radial diffusion, **↓** kurtosis | **Superior Longitudinal Fasciculus (SLF)** [123, 92, 108]  [**↓** FA, ↑ RD, ↑ MD, **↓** AK, ↓ RK, **↓** MK]   - **↓** anisotropy, ↑ radial diffusion, **↓** kurtosis   **Inferior Longitudinal Fasciculus (ILF)** [133, 111, 53]  [**↓** FA, ↑ RD, ↑ MD, **↓** AK, ↓ RK, **↓** MK]   - **↓** anisotropy, ↑ radial diffusion, **↓** kurtosis   **Inferior Fronto-Occipital Fasciculus (IFOF)** [127, 108, 64]  [**↓** FA, ↑ RD, ↑ MD, **↓** AK, ↓ RK, **↓** MK]   - **↓** anisotropy, ↑ radial diffusion, **↓** kurtosis   **Forceps Minor** [89, 152, 84] [*Medial*]  [**↓** FA, ↑AD, ↑ RD, ↑ MD, **↓** AK, ↓ RK, **↓** MK]   - **↓** anisotropy, ↑ diffusion, **↓** kurtosis   **Forceps Major** [89, 89, 84] [*Medial*]  [↑ AD, ↓ MK]   - ↑ axial diffusion, **↓** kurtosis   → [*Lateral* [R]] [68, 78, 84]  [↑ AD, ↑ RD, ↑ MD, **↓** AK, **↓** RK, **↓** MK]   - ↑ diffusion, **↓** kurtosis     **Anterior Thalamic Radiation** [106, 131, 81]  [↑ AD, ↑ RD, ↑ MD, **↓** AK, **↓** RK, **↓** MK]   - ↑ diffusion, **↓** kurtosis |

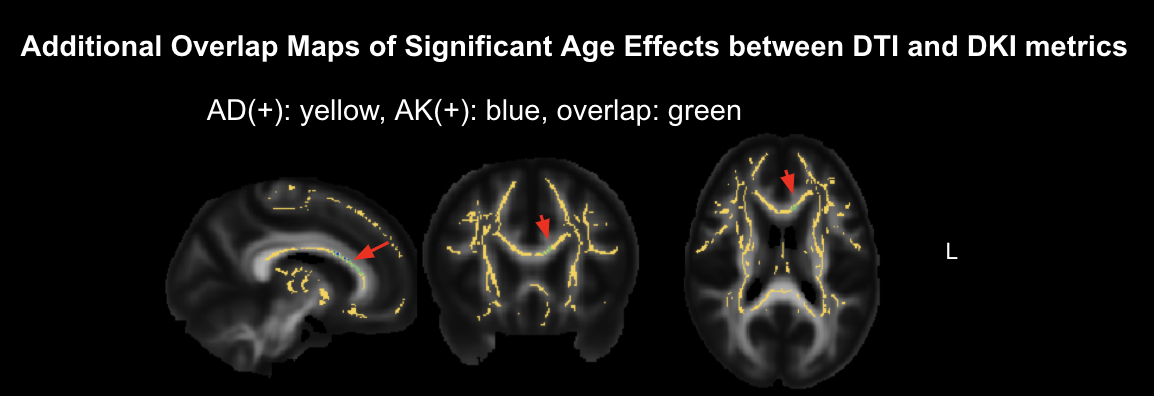

**Figure S1. An in-depth look at the positive AD and AK age associations in the anterior corpus callosum.** The overlap between AD and AK age effects is indicated by the red arrow. ‘L’ indicates neurological left.
